# Gas flow permeability and gas pressures in stilt roots of Red Mangrove (*Rhizophora mangle* L.)

**DOI:** 10.1101/2021.04.15.440060

**Authors:** Dennis G. Searcy

## Abstract

The Red Mangrove (*Rhizophora mangle*) is typically rooted in anoxic mud conditions that require special adaptations for root oxygenation. Each plant has multiple “stilt roots” that descend from upper branches and end in roots buried the mud. In cross section each stilt root consists of a core of porous aerenchyma surrounded by an impermeable layer of xylem, and outside the xylem there is a second layer aerenchyma. Oxygen must be provided to the mud roots through the aerenchyma either by diffusion or by gas flow, where the separate layers could provide up- and down-flow pathways. To test whether the stilt root’s properties were consistent with gas flow, conductivities were measured. A technique was developed that measured flow conductivities in S.I. scientific units without using a flow meter and a calibrated pressure gauge. The core aerenchyma was more permeable to gas flow than any other plant tissue, except for those stems that are hollow tubes. Because there was little lateral leakage from the core aerenchyma, it had pipe-like properties. In the outer aerenchyma gas conductivity was high, and gas flowed easily through large lenticels on the surface of the stilt root. To complete the calculations, gas pressures in the stilt roots were measured. The calculated gas flow rates through mud roots was not sufficient to supply O_2_ for root respiration, suggesting that diffusion may be the more important mechanism for these plants.

## Introduction

“Mangroves” is a term applied to the assemblages of small trees and shrubs that grow along shorelines in the tropics and in the subtropics (Mumby, Edwards et al. 2004). They provide a number of ecological services, but are threatened by human developments (*e.g*.- shrimp farms) and by sea level rise. When ocean levels rise, it is predicted that many mangroves will die by underwater root suffocation (Saintilan, Khan et al. 2020). Some mangroves live close to the maximum water depth in which they can survive, and so there is interest in the mechanism(s) by which O_2_ may be provided to their roots.

Oxygen can be provided to roots by the following routes. Diffusion through the soil is sufficient for typical upland plants. But when soil is saturated with water the rate of O_2_ diffusion is reduced by about 10,000 times (Beckett, Armstrong et al. 1988). There is some transport of dissolved O_2_ in sap flow (Hook, Brown et al. 1972), but diffusion of O_2_ down through the stems is sufficient for many smaller plants such as rice (Teal and Kanwisher 1966; Hook and Scholtens 1978; Beckett, Armstrong et al. 1988; Laan, Berrevoets et al. 1989; Allen 1997; Groot, van Bodegom et al. 2005). The distance limit for diffusion is about 25 cm (Laan, Berrevoets et al. 1989; Brune, Frenzel et al. 2000). When distances are much greater than 25 cm then gas flow through the stems is the likely mechanism of root oxygenation (Grosse, Büchel et al. 1998).

The importance of root aeration is well illustrated by the examples of water lilies and lotuses, both being rooted in deep water (Dacey 1980; 1981; Mevi-Shütz and Grosse 1988). Oxygen is supplied to the roots by convective gas flow through hollow stems (actually petioles). The air is pressurized initially by gas osmosis into the leaves, and typically separate petioles are used for up- and down-gas flows.

An informative exception is the Black Mangrove (*Avicennia marina*), which does not use gas flow but aerates its roots by diffusion. It grows on intertidal mud flats. There are finger-like projections (“pneumatophores”) that grow up to the air from the buried roots. Diffusion distances are short from the atmosphere to the buried roots. (Scholander, van Dam et al. 1955; Skelton and Allaway 1996).

The purpose of the present study was to measure the permeability to gas flow in stilt roots of *Rhizophora mangle* L. In the Western Hemisphere *R. mangle* is the plant that grows lowest along the shoreline, often in continuous water about 1 meter deep (Hogarth 2007). Adaptations include “stilt roots”, which are adventitious roots that grow down from upper branches into the water and terminate in “mud roots” in anoxic mud. (Fig. 1) The internal anatomy of stilt roots is unusual. See Chapman (1976), Tomlinson (1986), and Fig. 2 below. In cross section there are 2 concentric layers of porous aerenchyma. Aerenchyma is a plant tissue characterized by large intracellular air spaces that can facilitate O_2_ transport either by diffusion or by convective gas movement (Brix, Sorrell et al. 1992; Jackson and Armstrong 1999). In each stilt root, the 2 layers of aerenchyma are separated by a layer of hard, dense xylem. The histology has been described by Evans et al. (2005). The lower surface of each stilt root is covered by numerous large lenticels that, close above the surface of the water, can cover 25% of the stilt root surface. The well developed aerenchyma and lenticels are presumed to be adaptations for root aeration.

**Fig 1.**
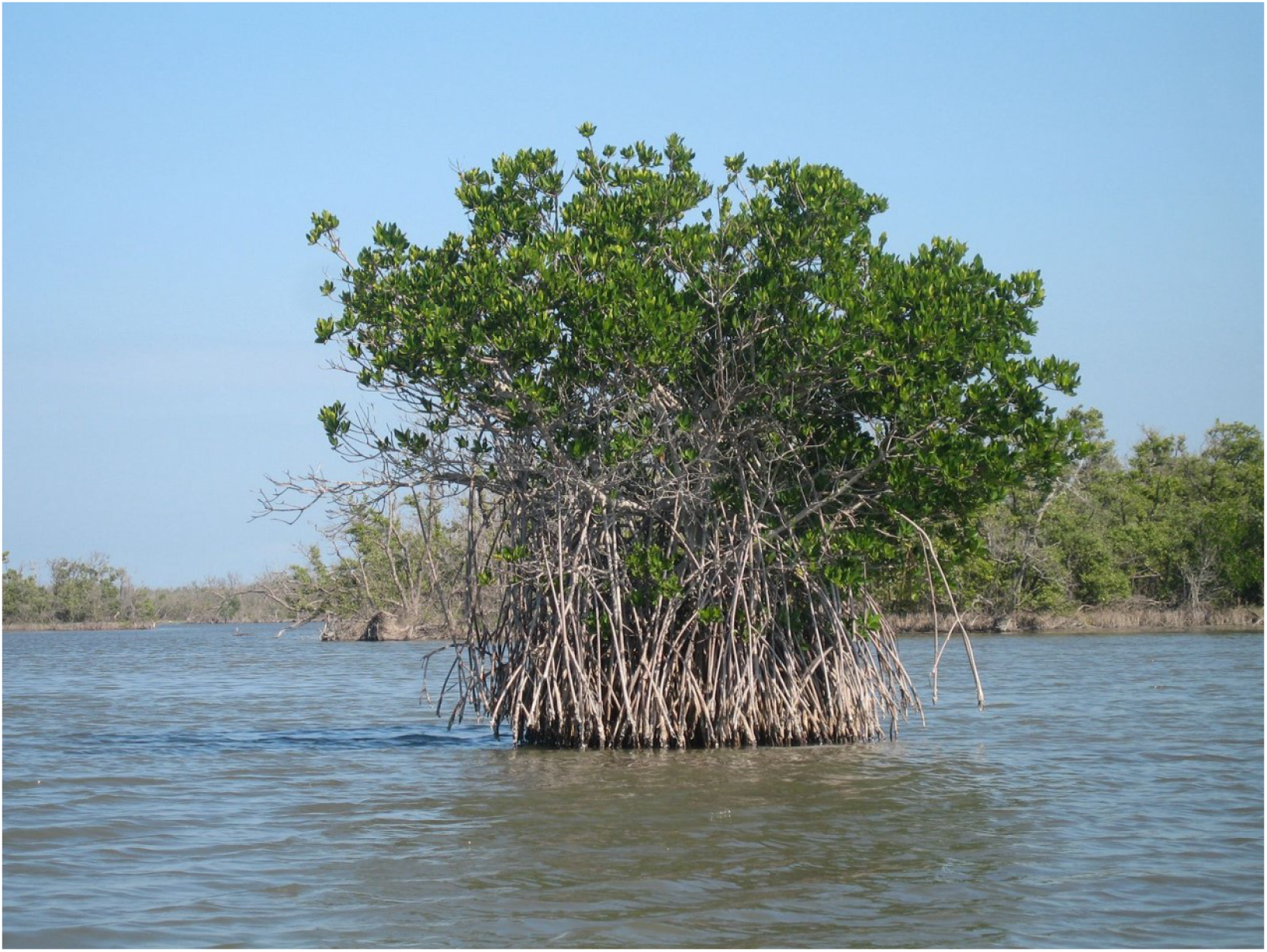
Red Mangrove (*Rhizophora mangle*). This plant is about 6 meters tall. Stilt roots drop from upper limbs into water about 1 meter deep. Below the water, the roots end in anoxic mud. Photograph by Andrew Tappert, Florida, 2007, and used under Gnu Free Documentation License. https://en.wikipedia.org/wiki/File:Red_mangrove-everglades_natl_park.jpg

**Fig 2.**
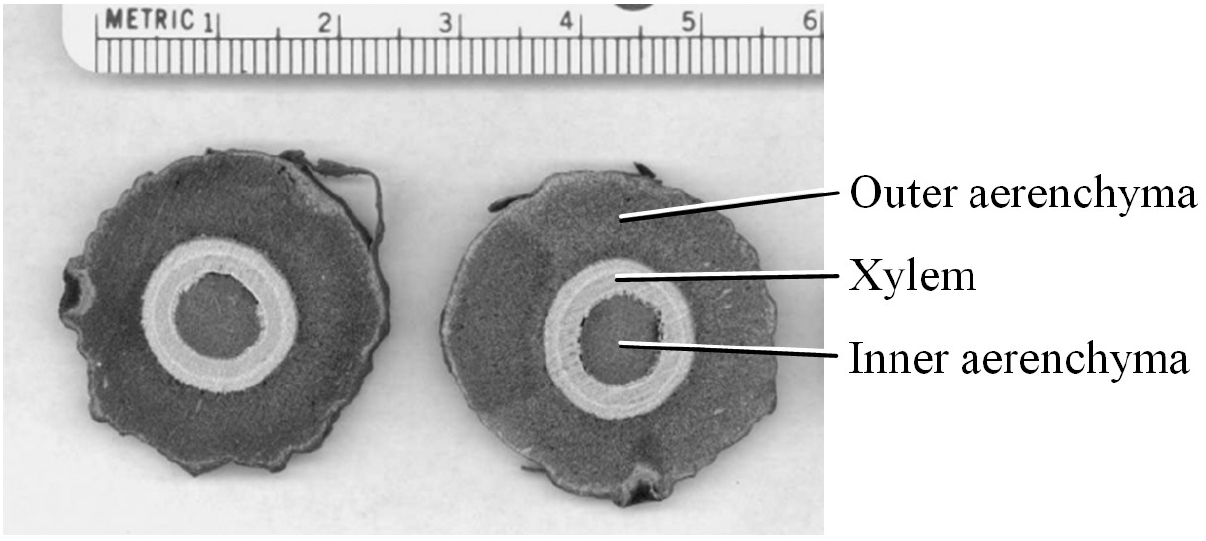
Cross-sections of a stilt root. The ruler is marked in cm.

Flow-through gas conductivity in *R. mangle* stilt roots was described by Scholander et al. (1955) who simply blew into one end of a section while holding the other end under water. Later Evans et al. (Evans, Okawa et al. 2005; Evans, de Leon et al. 2008; Evans, Testo et al. 2009) proposed a more detailed hypothesis for how *R. mangle* might provide O_2_ to its mud roots. Air is taken in and pressurized by enigmatic small “cork warts” on the undersides of leaves. Gas then flows down through limbs and prop roots to the buried mud roots. Since through-through aeration requires both up- and down-pathways, air flows up in outer aerenchyma and escapes into the atmosphere through the lenticels. Evans et al. tested their conjecture by using compressed air and by observing bubbles emitted from ends of sections held under water. The gas pressures were high and the flow data were not quantitative. The purpose of the present study was to obtain quantitative data using pressures that were more nearly natural.

Gas flow permeability in wood has been measured previously, for example to estimate dryness in lumber (Resch and Ecklund 1964; Sebastian, W. A. Côté et al. 1965; Petty 1970). Later, gas permeability was measured in aquatic and wetland plants, and so some technology exists. Instrumentation has included a source of pressurized air and electronic flow meters (Brix, Sorrell et al. 1992) (Armstrong, Armstrong et al. 1988; Evans, Testo et al. 2009).

In the present study, electronic instruments were used initially, but while working in ~0.5 m sea water sea water was accidentally splashed onto the instruments. They stopped working. Consequently a technique was improvised for measuring gas-flow permeability without electronic instrumentation.

The question addressed below is, are the properties of *R. mangle* stilt roots consistent with flow-through ventilation of the buried mud roots?

## Materials and methods

### Study sites

Cuttings of *R. mangle, Laguncularia racemosa,* and *Leucaena leucocephala* were obtained near Little Lameshur Bay, St. John, U.S. Virgin Islands. Collection permits were obtained from Virgin Islands National Park. For electronic measurements, specimens were taken to the nearby Virgin Islands Environmental Resource Station. (2) *Acer saccharum* was collected on the Univ. of Massachusetts, Amherst.

### Stilt root, branch, and stem cuttings

Stilt roots and branches without blemishes ~2.5 cm dia. were cut from plants, kept in shade, and used within 3 hours. Immediately before being measured each section was trimmed to 35 cm length.

Each end of a section of stilt root or branch was connected to short lengths of PVC tubing, heated and stretched and made air-tight with rubber cement. The PVC was connected to an electron pressure meter using 1.5 mm i.d. thick walled polyethylene tubing. Seals were tested by holding the connections under water and pressurized to 1 atm while watching for bubbles. When an experiment required a dry specimen the seals were water-tested after the experiment.

### Gas conductivity measurements

Fig. 3 shows the arrangement used to measure gas conductivity through a section of stilt root. During laboratory measurements an electronic pressure gauge and data logger were used. (Vernier Software & Technology, Beaverton, Oregon, USA).

**Figure 3.**
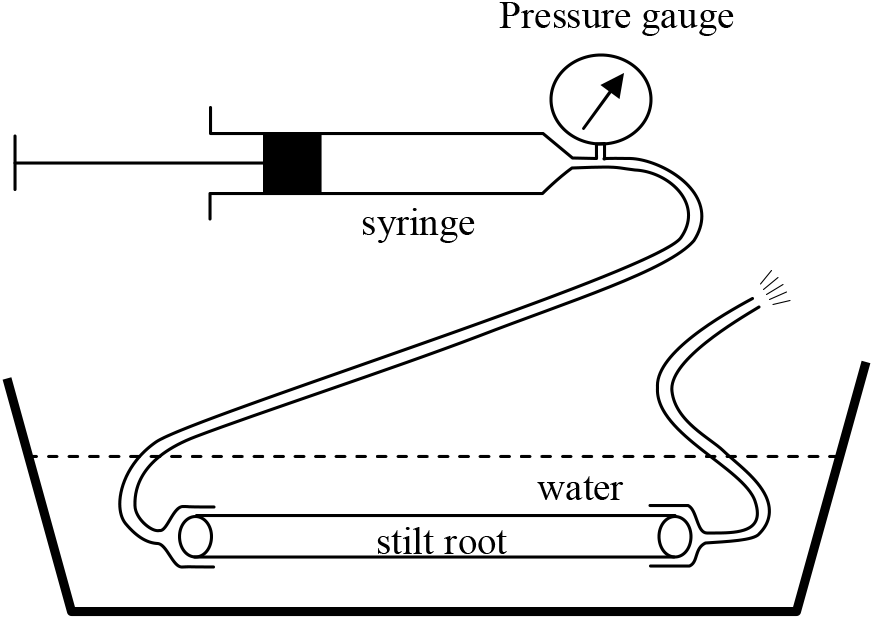
Laboratory arrangement for measuring gas conductivity. Tubing was attached to each end of a section of stilt root. A 60 cm^3^ syringe and electronic pressure gauge were connected to one end. The other end connected through tubing to the atmosphere. To make a measurement, the syringe plunger was pulled back from 55 cm^3^ to 60 cm^3^ creating ~0.1 atm relative vacuum in the syringe. As air flowed in through the stilt root, the pressure returned slowly to atmospheric. Pressures were recorded at 0.5 sec intervals.

As required by different experiments, sections of stilt roots were tested dry or wet, with the distal end plugged or open the atmosphere, and with positive or negative pressures (relative to atmospheric),

#### Measurements in the field using non-electronic instruments

A manometer was fashioned from 1.5 mm i.d. polyethylene tubing filled halfway with colored ethanol. When pressures made by the syringe caused a 14 cm difference between the arms of the manometer that corresponded to 1194 Pa = 0.12 atm. After pressurization gas flowed through the stem returning the pressure to atmospheric as reported on the manometer. Times for imposed gas pressures to return ½ and ¾ to atmospheric were measured using a stop watch.

### Effective gas volume inside the apparatus

For calculations it was essential to know the total gas space inside the apparatus including the syringe, tubing, and pressure gauge. That was measured by weighing the apparatus dry, filling it with water, and weighing it again.

### Gas conductivity calculations

When pressures were measured electronically gas conductivity coefficients were calculated using the following formula.

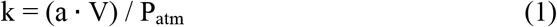

where “**a”** is the slope of a plot of ln |(P-P_atm_)/P| *vs.* time, and “V” is the volume of space inside the apparatus in cm^3^.

When pressures were measured using a manometer the following calculation was used:

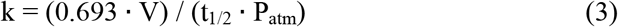

where **t**_½_ is the time in seconds required for pressure half-equilibration. Derivations of these formulas are in the Appendix.

### Gas pressures in stilt roots *in situ*

Pressures in the inner aerenchyma were recorded in stilt roots still attached to intact plants. A hole was drilled into a stilt root 1 m above the water or mud and connected to an electronic pressure gauge using small bore polyethylene tubing. The pressure gauge was connected to a digital data logger that recorded the pressure each minute. A second pressure gauge recorded simultaneously the ambient atmospheric pressure. After one day the two pressure gauges were reversed (to cancel out any slight mismatch between the gauges) and the recording continued for another day.

### O_2_ consumption by mud roots

A constant-pressure respirometer was improvised similar to that described by Gilson (Gilson 1963). Paper wetted with 5 M NaOH to absorb CO_2_. In operation, a syringe injected a measured amount of air in order to maintain constant pressure in the specimen chamber. Temperature was ambient: 29 C

Mud roots were removed from the mud, washed, and sorted. Only white, apparently healthy roots were used. They were cleaned again with a brush and sorted by size into rootlets > 1 mm and < 1 mm dia., and cut into 1 cm sections before being placed into to the respirometer chamber.

Oxygen consumption *in situ* (= “in *mudo*”) of buried mud roots was measured during low tide by cutting the stilt root 15 cm above the mud. A respiratory chamber (above) was modified by cutting hole in the bottom of the chamber and inserting it over the stump including both inner and outer layers of aerenchyma in the chamber.

The connection was stabilized and sealed with generous application of duct tape and then buried in mud to decrease the chance of gas leakage. Air was injected from a syringe so as to maintain constant pressure in the apparatus.

## Results

Typical gas flow data are shown in Fig. 4. In this example the plunger was pushed in by 5 cm^3^, creating inside a relative pressure of 10 KPa (0.1 atm). As air flowed in through the stilt root the pressure returned to atmospheric in an exponential curve which confirmed by the linearity of the logarithmic plot (Fig. 4B). These measurements used a dry stilt root section; air flowed in and out equally, which was not the case when wetted stilt roots were tested. (See later.)

**Figure 4.**
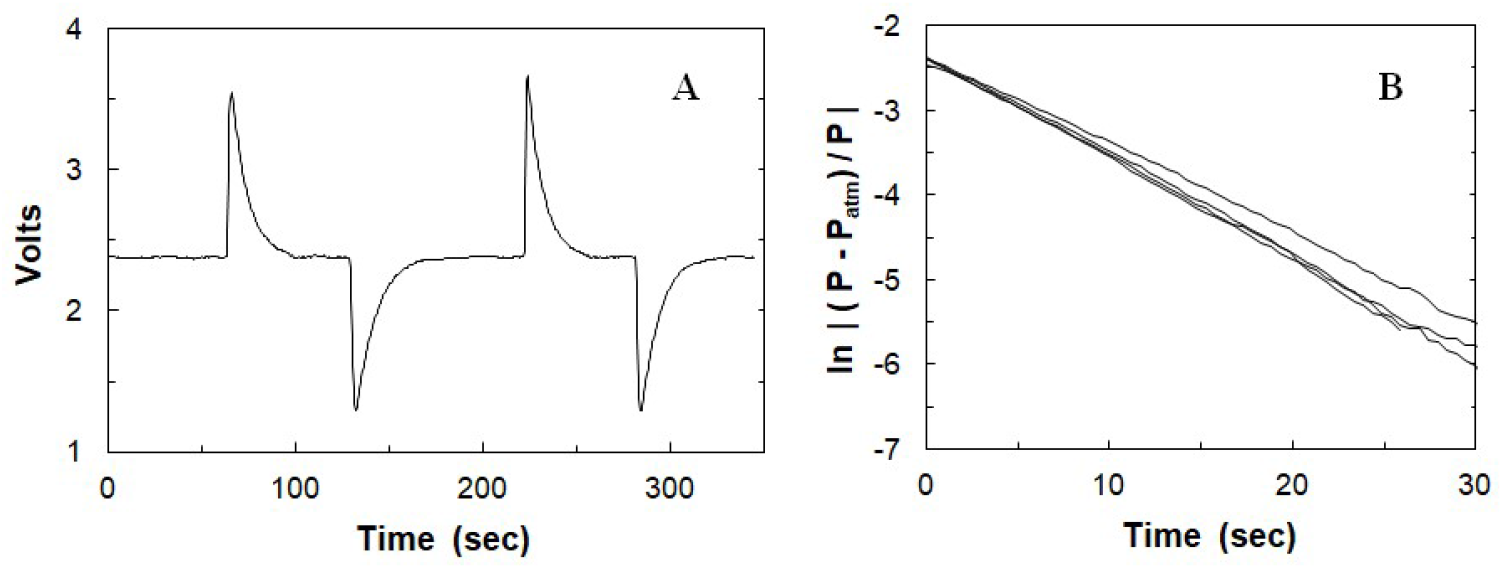
Typical gas flow data. Data were collected using a dry section of stilt root. The distal end was not plugged. First, the pressure inside the syringe and stilt root were allowed to equilibrate with atmospheric. Next, the syringe plunger was pushed in causing an increase in pressure and upward spike in voltage from the electronic pressure gauge. Then as air flowed out through the root, the pressure returned slowly to atmospheric. Next, the plunger was pulled out creating a relative vacuum. The manipulations were repeated. (**A**) Data as received in volts from the electronic pressure gauge. (**B**) The same data were converted to pressure units (Pa) and plotted as the logarithmic function shown. For display in Figure B the time axis was reset to zero at the start of each measurement. Data were converted from volts to Pa by a simple linear equation.

With tubing connected to one end of a stilt root section and the distal end plugged, air can escape through only the lateral surfaces. That was measured using dry root sections. See Table 1, rate of “Lateral gas escape.”) When the section was held under water and pressurized, streams of bubbles were observed escaping from the lenticels, confirming that lenticels were the route of gas flow through the skin. As the internal pressure declined, bubbles stopped streaming from some lenticels and then from others suggesting that a critical minimum pressure required to maintain gas flow through wet lenticels. Such flow was not simply proportional to pressure and the decline in gas flow was not exponential. Logarithmic plots were not linear. (Fig. 5)

**Table 1.** Lengthwise gas conductivity and lateral leakage measured in sections of stilt roots and woody branches. Gas Conductivities were measured using 35 cm sections of stilt roots and of branches, each ~2.5 cm dia. “Normalized Gas Conductivity” has been normalized for the dimensions of each structure by dividing by the cross-sectional area and multiplying by the length. “Lateral Escape” is gas conductivity through the surface measured using dry sections with their distal ends capped. Data are mean ± SE. The number of specimens measured is in the left column.

| *Species*<br>(number measured) | Gas Flow<br>Conductivity<br>(cm<sup>3</sup> s<sup>-1</sup> Pa<sup>-1</sup> × 10<sup>6</sup>) | Gas Conductivity<br>Normalized for<br>diameter and<br>length<br>(cm<sup>2</sup> s<sup>-1</sup> Pa<sup>-1</sup> × 10<sup>6</sup>) | Lateral Gas<br>Escape (cm<sup>3</sup> s<sup>-1</sup><br>Pa<sup>-1</sup> × 10<sup>6</sup>) |
| --- | --- | --- | --- |
| ***Rhizophora mangle* (Red Mangrove)** |  |  |  |
| Stilt root section |  |  |  |
| Inner aerenchyma (19) | 34 ± 9 | 1410 ± 320 | 1.5 ± 0.6 |
| Inner and outer<br>aerenchymas (9) | 127 ± 20 | 1220 ± 160 | 107 ± 15 |
| Upper branch section |  |  |  |
| Entire cut end (6) | 4.4 ± 0.6 | 37 ± 6 | 3.9 ± 0.9 |
| ***Laguncularia racemosa* (White Mangrove)** |  |  |  |
| Upper branch section (7) | 10 ± 1.7 | 76 ± 9 | 0.08 ± 0.6 |
| ***Leucaena leucocephala* (Lead Tree)** |  |  |  |
| Upper branch section(4) | 13 ± 1 | 134 ± 14 | 0.15 ± 0.2 |
| ***Acer saccharum* (Sugar Maple)** |  |  |  |
| Upper branch section (4) | 0.04 ± 0.16 | 0.18 ± 0.87 | 1.0 ± 0.7 |

**Figure 5.**
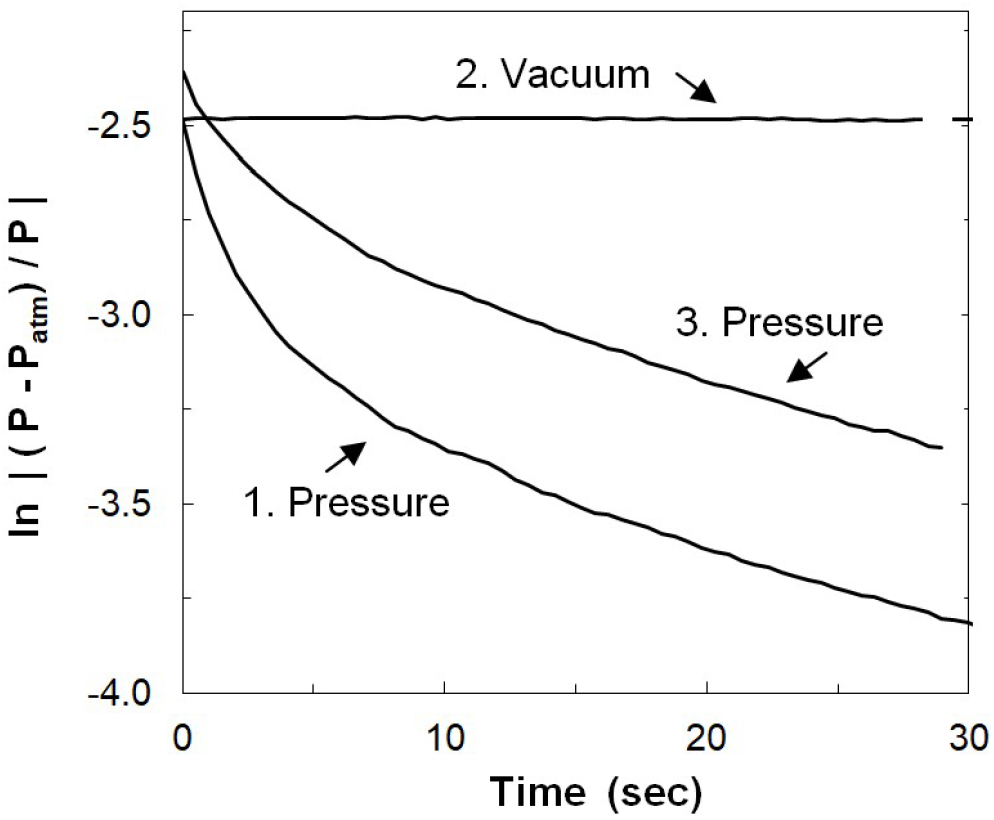
Gas flow into and out of the surface of a stilt root held under water. The distal end of the root section was plugged and then subjected to pressures of (1) +10 KPa, then (2) 10 KPa, and (3) +10 KPa. When pressurized, gas bubbled from the lenticels, and as the pressure decreased the gas flow declined but not exponentially. When a vacuum was applied there was no evidence of inward gas flow.

At the completion of the under water experiment the stilt root was dissected and was dry inside.

In order to test different pathways of gas flow the tubing was connected to the stilt root sections in different various patterns. For example, the distal end was plugged or not, and the root section was tested either dry, or wetted, or entirely under water. For example, in order to measure lengthwise gas flow down the length of the root section the distal end of a stilt root section was connected through tubing to the atmosphere while the section was held under water. (That is arrangement shown in Fig. 3.) When negative pressure was applied, gas flowed in but could enter only through the distal end of the section, and then it flowed the length of the section through both the inner and outer layers of aerenchyma.

To measure gas flow in the inner layer of aerenchyma by itself, the outer layer of aerenchyma was peeled back and tubing attached to the xylem layer. The data from these various arrangements of tubing and water are summarized in Table 1.

With the distal end capped, the only pathway for gas flow was through lenticels. With a dry stilt root section, gas flowed inward and outward equally well. But when the section was wetted and not under water, then gas flowed outward more easily than inward. (Fig. 6) The difference between flow rates in- and out-was significant by Student’s *t*-test: *P* < 0.01.

**Figure 6.**
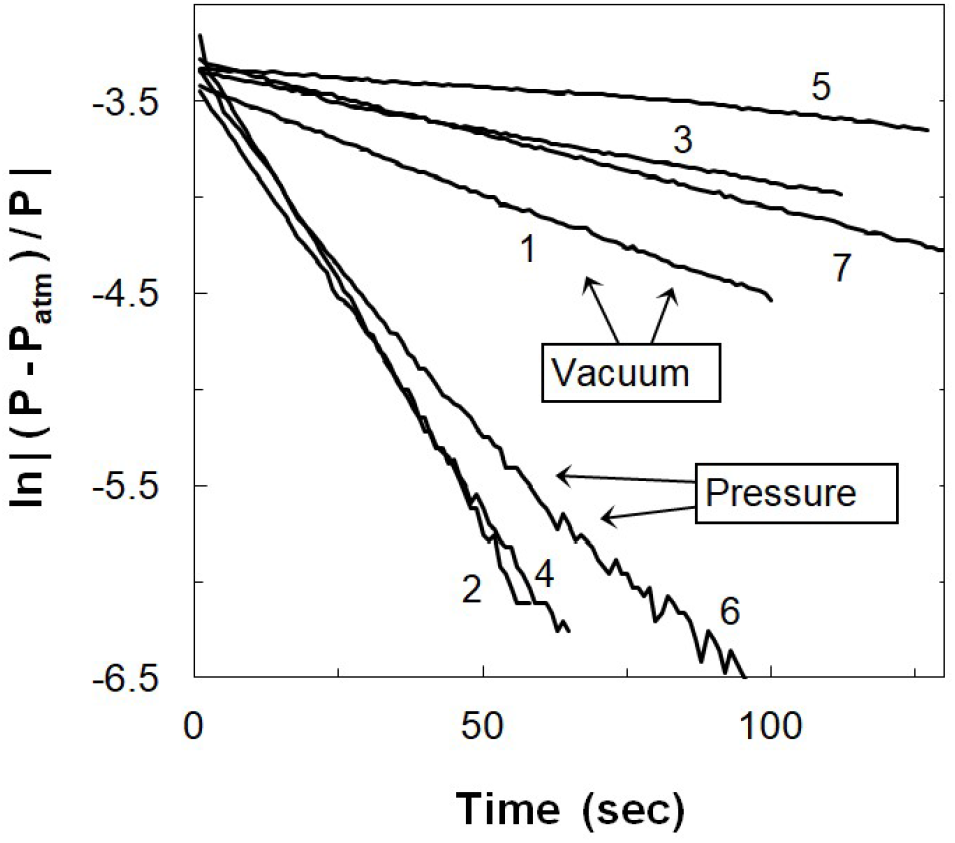
Gas flow into and out of a stilt root. The root section was wetted but not under water. The proximal end of the root section was attached to a pressure meter and a syringe; the distal end was capped. A single stilt root section was used for a series of experiments; the numbers in the figure indicate the sequence of measurements. Between each measurement the stilt root section was wetted again.

Table 1 includes gas conductivities measured in stems of several other species. *R. mangle* stilt roots had the highest gas conductivity of any tissue measured, which was over 3000 times greater than that through a branch of a typical New England forest tree. Gas conductivity was about 100 to 300 time higher in the stilt roots than in upper branches of *R. mangle* and upper braches of other mangrove species growing nearby.

### Mud root gas conductivity measured *in situ*

The hypothesis was that gas might flow down the inner aerenchyma, through the mud root, and up the outer aerenchyma. To test gas flow through mud roots, individuals were cut just above the water surface and the outer aerenchyma peeled back 1 cm. A syringe and manometer were attached to the inner aerenchyma and used to measure gas flow-through. (Table 2) At the top of the cut root, the outer aerenchyma connected with the atmosphere. Negative pressures to the inner aerenchyma were used to preclude the possibility of gas leakage out into the mud. The rates of pressure equilibration was measured by half-times using a manometer.

**Table 2.** Gas flow through buried roots. *R. mangle* stilt roots were cut 15 cm above the mud or water and the outer aerenchyma peeled back 1 cm. The inner aerenchyma was attached to 1 mm i.d. tubing connected to a 5 cm^3^ syringe and a manometer. *A. saccharum* (Sugar Maple) stems were similarly cut 15 cm above the soil and tubing attached to the entire cut end. Each section was ~2.5 cm dia. The *R. mangle* data represent 15 measurements on 3 different roots. The *A. saccharum* data include 9 measurements on 3 roots. Data are ± SE, N=3.

| *Rhizophora mangle*,<br>buried mud root | Mean ± SD |
| --- | --- |
| Time to $\frac{1}{2}$ pressure equilibration | 59 ± 5 sec |
| Time from $\frac{1}{2}$ equilibration to $\frac{3}{4}$ equilibration | 62 ± 7 sec |
| Gas volume in the apparatus | 20.4 ± 4 cm<sup>3</sup> |
| Pneumatic Conductivity $\times 10^6$ | 2.3 ± 0.1 cm<sup>3</sup> s<sup>-1</sup> Pa<sup>-1</sup> |
| *Acer saccharum*, root in soil |  |
| *Acer saccharum*, root in soil<br>Pneumatic Conductivity $\times 10^6$ | 0.008 ± 0.003 cm<sup>3</sup> s<sup>-1</sup> Pa<sup>-1</sup> |

Successive times to half equilibration and from half to three quarters equilibration were similar consistent with the mathematical model. (See Appendix.)

### Gas pressures in intact plants

Fig. 7 shows pressure recordings from the inner aerenchyma of a stilt root. The stilt root remained attached to its parent plant and mud root. Pressures are shown relative to ambient atmospheric pressure measured at the same time. Gas pressure in the inner aerenchyma became positive at dawn and remained positive for about 6 hours. Then the pressure became negative for the next 6 h hours. At night the pressure was weakly negative. The plants received morning sun but in the afternoon were shaded by higher trees behind them. These mangroves were at the edge of a salt pond where there were no oceanic tides.

**Fig. 7.**
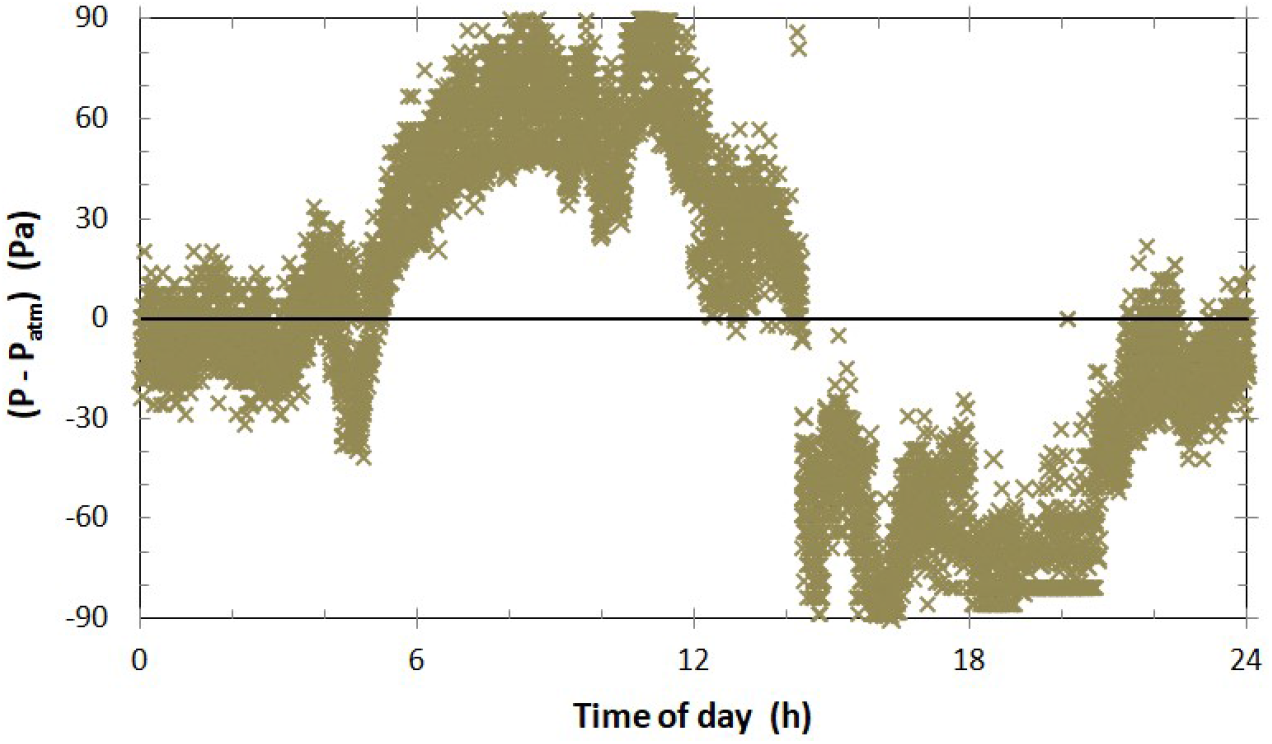
Pressures measured in the inner aerenchyma of a stilt root *in situ*. A hole was drilled into the inner aerenchyma and connected through tubing to a pressure gauge and recorder. A second gauge measured atmospheric pressure. The Y-axis shows the pressure difference between the inner aerenchyma and atmosphere. The figure summarizes nearly 30,000 paired measurements taken in 2 separate years and for a total of 8 days.

There was substantial variation in the pressure recordings that was of environmental origin. For example, when there was a rain shower the pressure dropped and when the sun reappeared the pressure went up.

### O_2_ consumption by mud roots

Oxygen consumption was measured also in excavated, cleaned rootlets ≤1 mm thickness. The rate of O_2_ consumption was 0.21 cm^3^ g^-1^ h^-1^ ±0.03 SE, N=5). For comparison the rate of O_2_ consumption of *Avicennia marina* roots (“black mangrove”) was 0.14 cm^3^ g^-1^ h^-1^ (Curran, Cole et al. 1986) and that of deepwater rice rootlets (*Oryza sativa*) was 0.54 cm^3^ g^-1^ h^-1^ (Guy 2003).

In order to estimate the weight of each mud root, 2 roots were dug out, cleaned and weighed. They weighed an average of 60 g. But some rootlets were embedded in compacted mud and could not be collected. But in each excavated mud root there was much dead and woody tissue. My guess is that each mud root was about 8 g metabolically active rootlets. Thus, O_2_ consumption for an 8 g of rootlets was 2 cm^3^ h^-1^.

A stilt root was cut 15 cm above the mud and a respirometer chamber fit to the exposed stump including inner and outer aerenchymas. The rate of O_2_ consumption was 2.7 cm^3^ h^-1^. That suggests that diffusion down the cut stump can be sufficient to supply mud root O_2_ consumption.

## Discussion

The utility of gas flow permeabilities reported above is they can be used to calculate gas flow rates: multiply the gas flow permeability by the pressure difference to obtain a gas flow rate. For example, if the permeability of the stilt root inner aerenchyma is 34 cm^3^ s^-1^ Pa^-1^×10^-6^ (Table 1) and the pressure difference is about 60 Pa (Fig. 7), the result is 7.3 cm^3^ h^-1^. The O_2_ consumption of mud roots was 2 cm^3^ h^-1^ suggesting that gas flow down the inner aerenchyma could supply sufficient O_2_ to a buried mud root.

But in the hypothetical path of gas flow the place that limits flow could be the mud root. The gas conductivity of mud roots was 2.3 cm^3^ s^-1^ Pa^-1^×10^-6^, and when multiplied by 60 Pa the calculated flow rate is 0.5 cm^3^ h^-1^. Furthermore, air is only 21% O_2_ so that the rate O_2_ supply is only 0.1 cm^3^ O_2_ h^-1^. That is only about 1/5 of the O_2_ that can be supplied by diffusion. So, in conclusion gas flow is unlikely to be important. The pipe-like properties of the inner aerenchyma are unexplained, as also is the function of gas flow pathway from leaves to mud roots described be Evans et al. (2005-2009), albeit the pathway does not have high sufficiently high gas conductivity for mud root oxygenation.

## Acknowledgments

Most of this work was done at the Virgin Islands Environmental Resource Station, St. John, USVI, while I accompanied a University of Massachusetts course in Tropical Field Biology.

## Appendix Calculation of gas conductivity

**Fig. 8.**
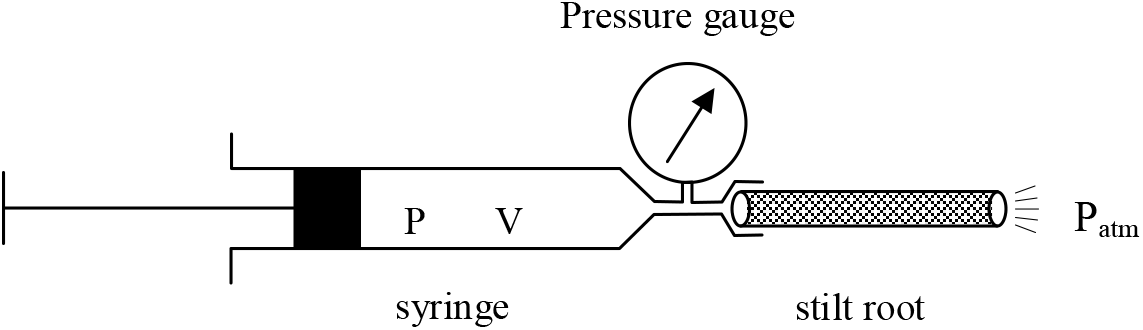
The diagram above shows a simplified experimental apparatus. “**P**” and “**V**” are the pressure and volume inside the syringe, including in the pressure gauge and tubing. **P_atm_** is atmospheric pressure.

Part of Darcy’s Law states that the rate of transport (gas or fluid) through a porous matrix is proportional to the pressure difference across the across the matrix (Leyton 1975). If the pressure applied is relative to ambient atmospheric pressure, that may be written:

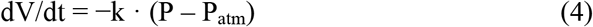

where **k** is the Gas Conductivity Constant. For a more complete discussion see Resch (1964) or Petty (1970).

Gases differ from liquids in that they are compressible. Volume is related to pressure by Boyle’s Law. One way to state that is:

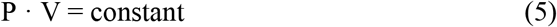

Differentiating and rearranging:

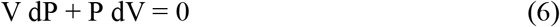

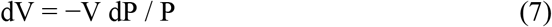

Substitute **Eqn. 9** into **Eqn. 6**, rearrange, and integrate. “C” is the constant of integration. Logarithms require positive values.

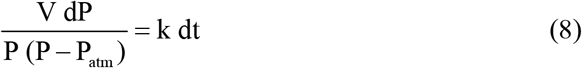

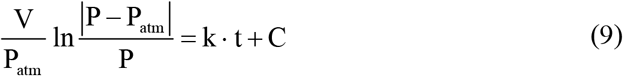

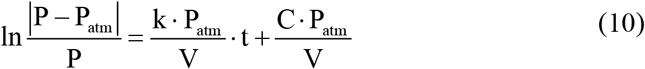

When one plots ln |(P – P_atm_)/P| vs. time the data should be a straight line with slope = k·P_atm_ /V. The Gas Conductivity (**k**) can be obtained by multiplying the slope by (V/P_atm_). When **V** is in units of cm^3^ and **P_atm_** is in Pa, then **k** will be in units of cm^3^ s^-1^ Pa^-1^.

A remarkable feature of the plot described above is that the result is in scientific S.I. units even though there is no flow meter and the pressure meter need not be calibrated: the y-axis is without units. Any pressure unit can be used. The rate of gas flow can be measured by either the slope of the plot or by the “half-time”. Halftimes are used when collecting non-electronic data such as in the field using a manometer.

The utility of Gas Conductivity is that actual gas flow through any object can by calculated by multiplying by the pressure differential (in Pa) measured across the object.

The technique can be adapted to a broad range of gas conductivities by changing **V** by changing the syringe size. One can use a small syringe to measure a slow gas flow rate or a large syringe for a rapid flow rate.

### Normalized Conductivity

To enable comparisons between objects of different sizes, “Normalized Gas Conductivity” can be used. A more complete statement of Darcy’s Law includes that the rate of pressurized flow through porous medium is proportional to cross-sectional area (A) in inversely proportional to distance (L) (Leyton 1975; Vogel 1994). Thus, Gas Conductivity measurements can be normalized for dimensions as follows.

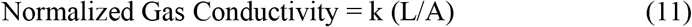

The units of Normalized Gas conductivity are: cm^3^ gas flow · s^-1^ · Pa^-1^ · cm length · (cm^2^ cross sectional area)^-1^. After cancellations that becomes cm^2^ s^-1^ Pa^-1^.

## References

Armstrong, J., W. Armstrong, et al. (1988). “*Phragmites australis:* a critical appraisal of the ventilating pressure concept and an analysis of resistance to pressurized gas flow and gaseous diffusion in horizontal rhizomes.” New Phytol. 110(3): 383–389.

Beckett, P. M., W. Armstrong, et al. (1988). “On the Relative Importance of Convective and Diffusive Gas-Flows in Plant Aeration.” New Phytol 110(4): 463–468.

Brix, H., B. K. Sorrell, et al. (1992). “Internal pressurization and convective gas flow in some emergent freshwater macrophytes.” Limnol. Oceanogr. 37(7): 1420–1433.

Brune, A., P. Frenzel, et al. (2000). “Life at the oxic-anoxic interface: microbial activities and adaptations.” Fems Microbiology Reviews 24(5): 691–710.

Chapman, V. J. (1976). Mangrove Vegetation. Vaduz, J. Cramer.

Curran, M., M. Cole, et al. (1986). “Root aeration and respiration in young mangrove plants (*Avicennia marina* (Forsk.) Vierh.).” J. Exp. Bot. 37: 1225–1233.

Dacey, J. W. H. (1980). “Internal winds in water lilies: an adaptation for life in anaerobic sediments.” Science 210: 1017–1019.

Dacey, J. W. H. (1981). “Pressurized ventilations in the yellow waterlily.” Ecology 62: 1137–1147.

Evans, L. S., M. F. de Leon, et al. (2008). “Anatomy and morphology of *Rhizophora stylosa* in relation to internal airflow and Attim’s plant architecture.” J. Torrey Bot. Soc. 135(1): 114–125.

Evans, L. S., Y. Okawa, et al. (2005). “Anatomy and morphology of red mangrove *(Rhizophora mangle)* plants in relation to internal airflow.” J. Torrey Bot. Soc. 132(4): 537–550.

Evans, L. S., Z. M. Testo, et al. (2009). “Characterization of internal airflow within tissues of mangrove species from Australia: leaf pressurization processes.” J. Torrey Bot. Soc. 136(1): 70–83.

Gilson, W. E. (1963). “Differential respirometer of simplified and improved design.” Science 141(3580): 531–532.

Groot, T. T., P. M. van Bodegom, et al. (2005). “Gas Transport through the Root--shoot Transition Zone of Rice Tillers.” Plant and Soil 277(1): 107–116.

Grosse, W., H. B. Büchel, et al. (1998). Root aeration in wetland trees and its ecophysiological significance. Coastally Restricted Forests. A. D. Laderman. New York, Oxford Univ. Press: 293–305.

Guy, J. D. K. (2003). “Rice root properties for internal aeration and efficient nutrient acquisition in submerged soil.” New Phytologist 159(1): 185–194.

Hogarth, P. J. (2007). The Biology of Mangroves and Seagrasses. Oxford, Oxford.

Hook, D. D., C. L. Brown, et al. (1972). “Aeration in trees.” Bot. Gaz. 133(4): 443–454.

Hook, D. D. and J. R. Scholtens (1978). Adaptations and flood tolerance of tree species. Plant Life in Anaerobic Environments. D. D. Hook and R. M. M. Crawford. Ann Arbor, Michigan, Ann Arbor Science Publishers: 299–331.

Jackson, M. B. and W. Armstrong (1999). “Formation of aerenchyma and the processes of plant ventilation in relation to soil flooding and submergence.” Plant Biology 1(3): 274–287.

L. H. Allen, J. (1997). “Mechanisms and rates of O2 transfer to and through submerged rhizomes and roots via aerenchyma.” Proc. Soil Crop Sci. Soc. Florida 56: 41–54.

Laan, P., M. J. Berrevoets, et al. (1989). “Root morphology and aerenchyma formation as indicators of the flood-tolerance of *Rumex species*.” J. Ecol. 77(3): 693–703.

Leyton, L. (1975). Fluid Behavior in Biological Systems. Oxford, Clarendon Press.

Mevi-Shütz, J. and W. Grosse (1988). “A two way gas transport system in *Nelumbo nucifera*.” Plant Cell Environ. 11: 27–34.

Mumby, P. J., A. J. Edwards, et al. (2004). “Mangroves enhance the biomass of coral reef fish communities in the Caribbean.” Nature 427(6974): 533–536.

Petty, J. A. (1970). “Permeability and structure of the wood of sitka spruce.” Proc. R. Soc. Lond. B Biol. Sci. 175(1039): 149–166.

Resch, H. and B. A. Ecklund (1964). “Permeability of wood exemplified by measurements on Redwood.” Forest Prod. J. 14(5): 199–206.

Saintilan, N., N. Khan, et al. (2020). “Thresholds of mangrove survival under rapid sea level rise.” Science 368: 1118–1121.

Scholander, P. F., L. van Dam, et al. (1955). “Gas exchange in the roots of mangroves.” Am. J. Bot. 42: 92–98.

Sebastian, L. P., J. W. A. Côté, et al. (1965). “Relationship of gas phase permeability to ultrastructure of White Spruce wood.” Forest Prod. J. 15(9): 394–404.

Skelton, N. J. and W. G. Allaway (1996). “Oxygen and pressure changes measured in situ during flooding in roots of the Grey Mangrove *Avicennia marina* (Forssk.) Vierh.” Aquat. Bot. 54(2-3): 165–175.

Teal, J. M. and J. W. Kanwisher (1966). “Gas transport in the marsh grass *Spartina alternifora*” J. Exp. Bot. 17: 355–361.

Tomlinson, P. B. (1986). The Botany of Mangroves. Cambridge, Cambridge Univ. Press.

Vogel, S. (1994). Life in Moving Fluids. Princeton, NJ, Princeton Univ. Press.

